# Pathogen diversity drives the evolution of promiscuous peptide binding of human MHC-II alleles

**DOI:** 10.1101/371054

**Authors:** Máté Manczinger, Gábor Boross, Lajos Kemény, Viktor Müller, Tobias L. Lenz, Balázs Papp, Csaba Pál

## Abstract

Major histocompatibility complex (MHC) molecules mediate the adaptive immune response against pathogens. Certain MHC alleles are generalists: they present an exceptionally large variety of antigenic peptides. However, the functional implications of such elevated epitope binding promiscuity in the MHC molecules are largely unknown. According to what we term the pathogen-driven promiscuity hypothesis, exposure to a broad range of pathogens favors the evolution of highly promiscuous MHC variants. Consistent with this hypothesis, we found that in pathogen-rich geographical regions, humans are more likely to carry promiscuous MHC class II DRB1 alleles, and the switch between high and low promiscuity levels has occurred repeatedly and in a rapid manner during human evolution. We also show that selection for promiscuous peptide binding shapes MHC genetic diversity. In sum, our study offers a conceptually novel mechanism to explain the global distribution of allelic variants of a key human immune gene by demonstrating that pathogen pressure maintains promiscuous MHC class II alleles. More generally, our work highlights the hitherto neglected role of epitope binding promiscuity in immune defense, with implications for medical genetics and epidemiology.

---

The major histocompatibility complex (MHC) genes in vertebrates encode cell-surface proteins and are essential components of adaptive immune recognition (1). MHC proteins are endowed with highly variable peptide-binding domains that bind short protein fragments. The MHC region is one of the most polymorphic gene clusters in vertebrate genomes (2). Pathogen-driven balancing selection (PDBS) is considered largely responsible for the observed exceptionally high levels of genetic diversity (3–5) (Fig. 1A). Balancing selection may act through heterozygote advantage or frequency-dependent selection favoring rare MHC alleles (6).

**Fig. 1.**
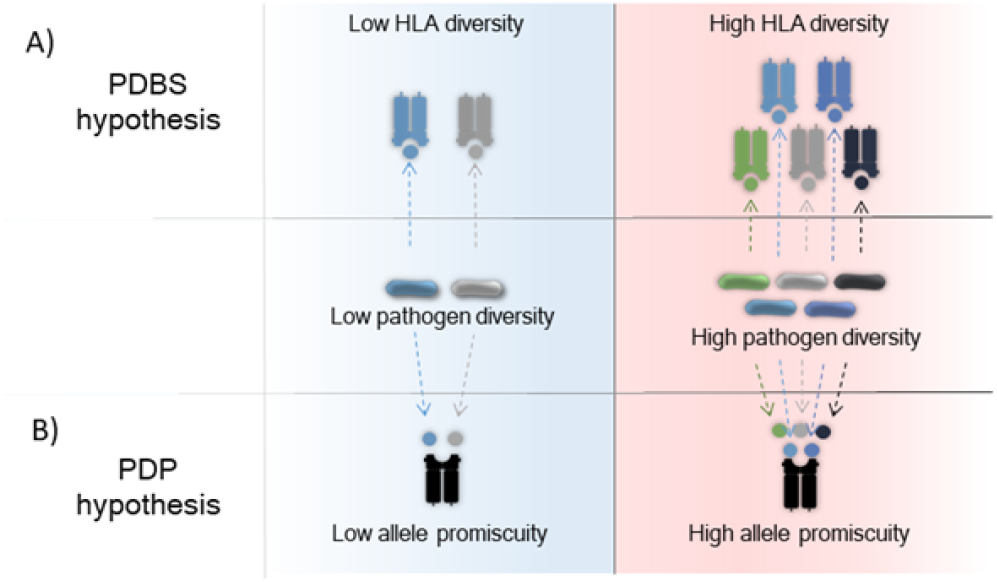
Allelic diversity and epitope binding promiscuity of HLA molecules jointly shape pathogen recognition. During the co-evolution of the human immune system and pathogens, HLA molecules evolved to recognize emerging pathogens. (**A**) HLA allelic diversity is maintained by pathogen-driven balancing selection (PDBS). (**B**) We propose that high pathogen diversity also selects for alleles with elevated epitope binding promiscuity in these areas (PDP: pathogen-driven promiscuity).

Multiple lines of evidence from humans and other vertebrates are consistent with the PDBS hypothesis (7). However, the PDBS hypothesis cannot account for a phenomenon that has only recently been appreciated in its full significance. There is a substantial variation in the size of the bound and presented antigen repertoire among different MHC variants. Certain MHC alleles appear to be promiscuous and are capable of binding an exceptionally large set of epitope peptide segments, with implications on immunocompetence and pathogen resistance (8).

According to what we here term the pathogen-driven promiscuity (PDP) hypothesis (Fig. 1B), selection should favor such promiscuous alleles in pathogen-rich geographical regions, as they promote immune response against a potentially broader range of pathogens (8). This selection process is analogous to PDBS-based processes in the way it is driven by pathogen diversity. However, while PDBS leads to a larger number of alleles in a given population, PDP hypothesis affects the binding properties of individual alleles. Therefore, PDBS and PDP represent two complementary ways in which MHC genes respond to pathogen-driven selection, with allelic diversity and increased epitope binding repertoire of individual MHC variants jointly shaping the recognition of foreign peptide segments (Fig. 1).

The proposed hypothesis (PDP) predicts that MHC promiscuity should provide protection against a broad range of pathogens at the individual level and at the same time shape the geographical distribution of MHC alleles. As a consequence, in regions of high pathogen diversity, human populations should carry promiscuous MHC alleles. Moreover, as migrating human populations have been exposed to changing sets of pathogens (9), shifts in MHC promiscuity level should have occurred repeatedly and in a rapid manner during the course of human evolution.

To test these predictions, we first focused on the human HLA class II DRB1 gene, for four reasons. First, DRB1 is the most variable HLA class II locus with over 2000 registered alleles (10). Together with HLA-DRA, HLA-DRB1 encodes the heterodimeric HLA-DR protein complex, but HLA-DRA is basically invariant. Second, DRB1 shows the strongest signature of selection among HLA class II loci (11), and it has diversified very rapidly in the human lineage (12). Many of these alleles appear to be human-specific and most likely evolved after the migration of ancestral human populations out of Africa (12). These periods have been associated with human populations encountering numerous new pathogens (9, 13). For other HLA class II loci, the level of genetic diversity is lower (10, 11), probably driven by evolutionary forces unrelated to pathogens (14). Third, epitope binding prediction algorithms show higher accuracy for DRB1, than for other HLA class II loci (15, 16). Finally, the abundance of DRB1 on the cell surface is especially high compared to other HLA class II receptors, indicating special significance of these molecules in antigen presentation (17).

As HLA-DR presents peptide epitopes derived mainly from extracellular proteins to T-cells (1), the pathogen-driven promiscuity hypothesis additionally predicts that primarily extracellular pathogen richness should influence the geographical distribution of promiscuous HLA-DRB1 alleles. Estimates on epitope binding promiscuity were derived from two sources: experimental assays that measured individual peptide-MHC interactions in vitro and systematic computational predictions. In a series of analyses, we show that predictions of the pathogen-driven promiscuity hypothesis are upheld, regardless of how HLA-DRB1 promiscuity level is estimated.

## Results

### Estimating HLA promiscuity level

Given that large-scale experimental assays to measure individual peptide-MHC interactions are extremely tedious, we first employed established bioinformatics tools to predict the binding affinities of experimentally verified epitope peptides for a panel of 160 nonsynonymous HLA-DRB1 alleles, all of which are present at detectable frequencies in at least one human population (18–20). The set of investigated epitopes was derived from the Immune Epitope Database (IEDB) and contains 2691 peptide epitopes of 71 pathogens known to be bound by certain HLA II variants (21) (Dataset S1). Epitopes showing high levels of amino acid similarity to each other were excluded from the analysis (See Methods). Most included epitopes are 15 to 20 amino acids long, and are found in only one of the 71 pathogens (SI Appendix, Fig. S1, Dataset S1). The NetMHCIIpan algorithm was used to predict individual epitope-MHC interactions (15), not least because it outperforms other prediction algorithms (16). The breadth of epitope binding repertoire or, shortly, the level of promiscuity of individual HLA-DRB1 alleles was estimated as the fraction of epitopes with a binding affinity stronger than 50 nM to the given MHC molecule. This threshold corresponds to high-affinity binding, which is frequently observed in MHC molecules displaying immunodominance (22). We found large variation in promiscuity levels across HLA-DRB1 alleles (SI Appendix, Fig. S2). Using a smaller dataset with information from both approaches, we show that the computationally predicted and the empirically estimated promiscuity values are strongly correlated with each other (Spearman’s rho: 0.78, P = 0.004, SI Appendix, Fig. S3). Moreover, our results are robust to changes in the affinity threshold (SI Appendix, Fig. S4, A to C), usage of other prediction algorithms (SI Appendix, Fig. 4D) and different epitope data sets (SI Appendix, Fig. 4E). As expected, promiscuous HLA-DRB1 alleles can present epitopes from a broader range of pathogen species (SI Appendix, Fig. S5). Reassuringly, there was no correlation between allele promiscuity values and the amount of data per allele used for the training of the algorithm (Spearman’s rho: −0.38, P = 0.21).

### Global distribution of promiscuous HLA alleles

Taking advantage of the confirmed reliability of computational predictions, we next investigated the geographic distribution of HLA-DRB1 alleles. We first collected high quality HLA-DRB1 allele prevalence data of 96 human populations residing in 43 countries from two databases and an article (18–20). The weighted average of promiscuity level in each population was calculated based on the promiscuity values and allele frequencies of individual alleles in the population (See Methods). The analysis revealed a large variation in mean promiscuity across geographical regions and the corresponding human populations (SI Appendix, Table S1). Importantly, several distantly related, but highly promiscuous alleles contribute to this pattern (SI Appendix, Table S1). Notably, an especially high allelic promiscuity level was found in South East Asia, an important hotspot of emerging infectious diseases (23). To minimize any potential confounding effect of high genetic relatedness between neighboring populations, we merged populations with similar HLA allele compositions for all further analyses (See Methods).

### Link between pathogen diversity and HLA promiscuity level

Using the Global Infectious Diseases and Epidemiology Network (GIDEON), we compiled a dataset on pathogen richness in the corresponding 43 geographic regions (24). It consists of 95 diseases caused by 168 extracellular pathogens, including diverse bacterial species, fungi, protozoa and helminthes. Using the same protocol, we additionally compiled a dataset on the prevalence of 149 diseases in the same regions caused by 214 viral and other obligate intracellular pathogens. The dataset and methodology employed for the analysis are standardized and have been used previously in similar contexts (7, 25).

We report a strong positive correlation between extra-cellular pathogen diversity and mean promiscuity: HLA-DRB1 alleles that can bind epitopes from a broader range of pathogens are more likely to be found in regions of elevated pathogen diversity (Fig. 2A). As mentioned above, the proposed hypothesis also predicts that HLA-DRB1 promiscuity should be specifically associated with extracellular, but not intracellular pathogen diversity. In line with this expectation, we found no significant association between HLA-DRB1 promiscuity level and diversity of intracellular pathogens (Fig. 2B). We conclude that the geographical distribution of promiscuous HLA-DRB1 alleles has been shaped by the diversity of extracellular pathogens.

**Fig. 2.**
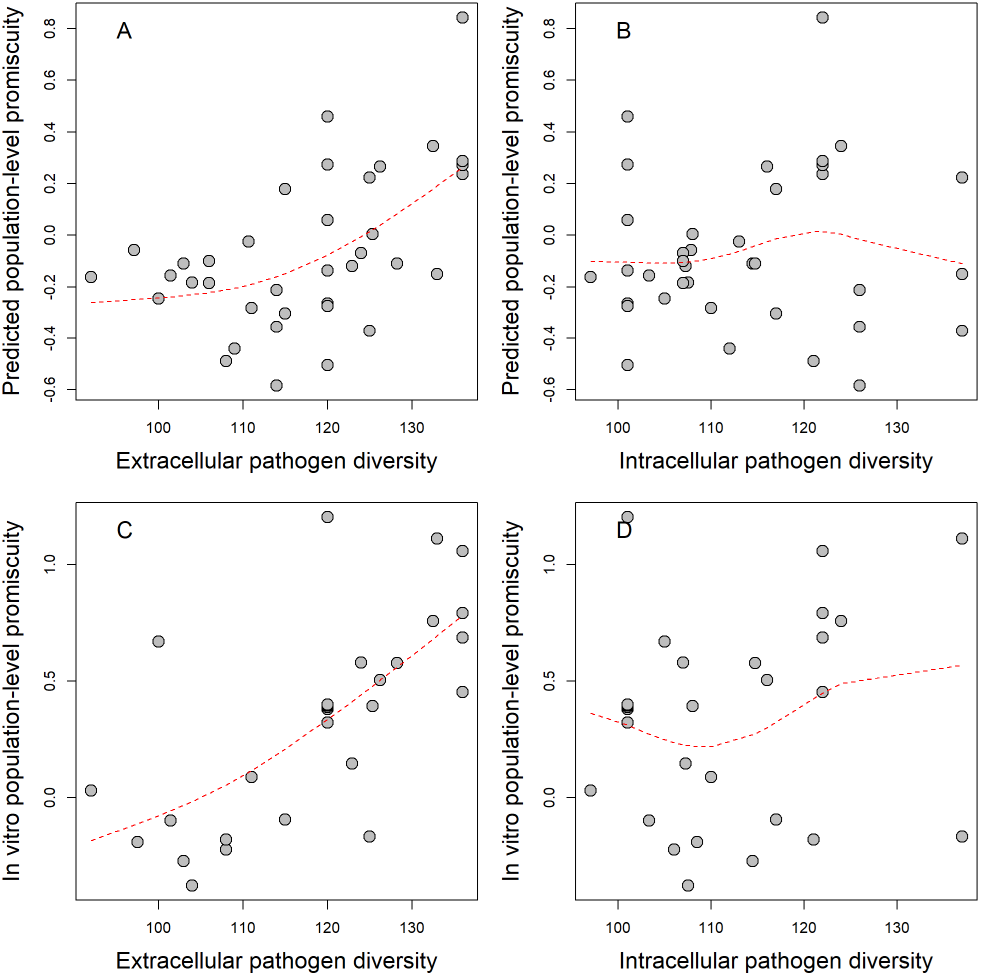
Relationship between epitope binding promiscuity and pathogen diversity. Normalized population-level promiscuity of HLA-DRB1 alleles is shown as the function of extracellular pathogen diversity as approximated by species count. Promiscuity scores were calculated based on standardized (i.e. z-score) allele promiscuity values and were averaged in each population group (see Methods). Significant correlations were found between extracellular pathogen species count and (**A**) predicted allele promiscuity level in 37 groups (Spearman’s rho: 0.5, P = 0.002) and (**C**) in vitro promiscuity level in 28 groups (Spearman’s rho: 0.7, P = 3 × 10^−5^). No significant correlation was found between intracellular pathogen diversity and (**B**) predicted and (**D**) in vitro promiscuity at the HLA-DRB1 locus (Spearman’s rho: 0.04 and 0.21, P = 0.81 and 0.29, respectively). Dashed lines indicate smooth curve fitted using cubic smoothing spline method in R (see Methods). Population groups were created using the 15th percentile genetic distance cutoff (see Methods). For results obtained upon using alternative distance cutoff values, see Supplementary Data 2.

The above results hold – and are even stronger – when estimates on promiscuity were derived from data of empirical in vitro binding affinity assays (shortly in vitro promiscuity), downloaded from the IEDB database (21) (Fig. 2, C and D, SI Appendix, Table S2 and Dataset S2). However, these results do not exclude the possibility that the geographical link between pathogen diversity and promiscuity is indirect. More direct support on the causal relationship between the two variables comes from analysis of prior human genetic studies. The data indicate that multiple allele groups with high promiscuity levels are associated with protection against a broad range of infectious diseases (SI Appendix, Table S3).

The data also indicate local adaptation towards elevated promiscuity under diverse pathogen pressure. The HLA-DRB1*12:02 allele is prevalent in specific regions of South East Asia. Compared to other alleles detected in this region, HLA-DRB1*12:02 has an exceptionally high promiscuity value (Fig. 3A). The high frequency of HLA-DRB1*12:02 has been previously suggested to reflect pathogen-driven selection during migration of a Mongolian population to South China (26). Indeed, this allele is associated with protection from several infectious diseases caused by extracellular pathogens (SI Appendix, Table S3), many of which are endemic in South East Asia (27, 28). Remarkably, the frequency of this allele increases with extracellular pathogen diversity in this region (Fig. 3B). Together, these observations support the hypothesis that promiscuous epitope binding of HLA-DRB1 alleles is favored by selection when extracellular pathogen diversity is high.

**Fig. 3.**
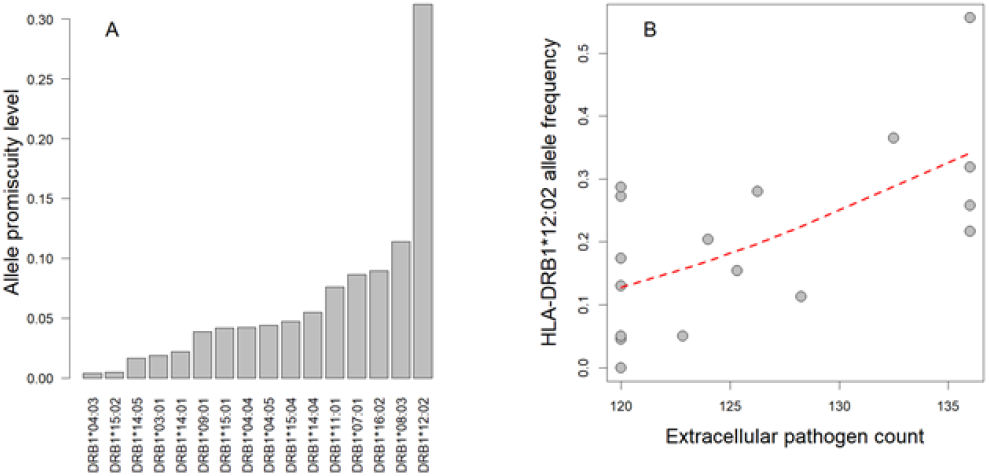
HLA-DRB1*12:02 allele promiscuity level and extracellular pathogen diversity in Southeast Asia. (**A**) HLA-DRB1*12:02 has exceptionally high promiscuity level compared to other alleles. The figure shows alleles with at least 10% frequency in at least one population in Southeast Asia. Predicted allele promiscuity values are shown. (**B**) The frequency of DRB1*12:02 increases with extracellular pathogen diversity across populations (Spearman’s rho: 0.57, P = 0.017). Populations resided in China, Japan, South Korea, Indonesia, Malaysia and the Philippines were included in the analysis. Red curve indicates smooth curve fitted using cubic smoothing spline method in R (see Methods).

### Evolution of promiscuous HLA alleles

An important unresolved issue is how promiscuity has changed during the course of human evolution. Under the assumption that local pathogen diversity drives the evolution of epitope recognition of HLA class II alleles, promiscuity as a molecular trait should have evolved rapidly as human populations expanded into new territories. To investigate this issue, we combined an established phylogeny of HLA-DRB1 alleles (29) with predicted epitope binding promiscuity values. We found that alleles with a high promiscuity level have a patchy distribution across the tree (SI Appendix, Fig. S6), indicating that high promiscuity has multiple independent origins. To investigate this observation further, we selected a set of 96 HLA-DRB1 alleles with a detectable frequency in at least one human population and appropriate sequence data (See Methods). A comparison of all pairs of these alleles revealed that even very closely related alleles show major differences in promiscuity levels (Fig. 4A). For example, alleles belonging to the HLA-DRB1*13 group show over 98% amino acid sequence identity to each other, but display as much as 57-fold variation in the predicted promiscuity levels. We conclude that the switch between high and low promiscuity levels has occurred repeatedly and in a rapid manner during the allelic diversification of the HLA-DRB1 locus.

**Fig. 4.**
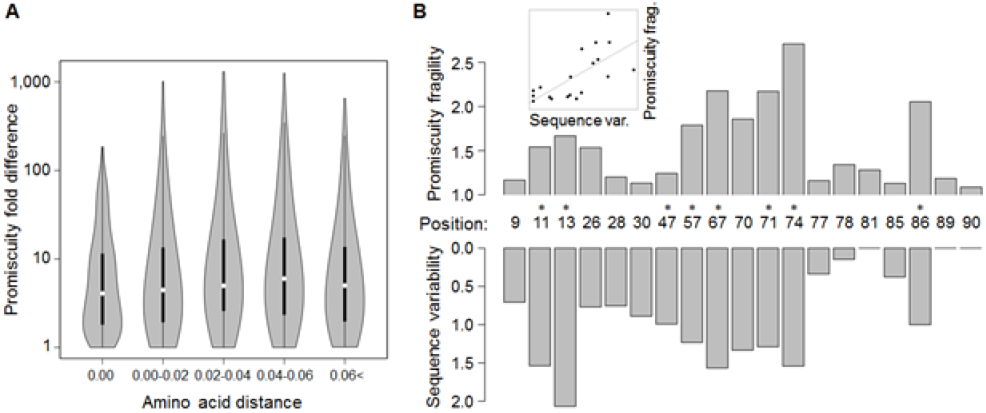
Promiscuity changes rapidly during evolution and might be a selectable trait. (**A**) For all pairs of selected alleles, predicted promiscuity difference between two HLA-DRB1 alleles is shown as a function of amino acid distance measured after excluding the epitope binding region. Large differences in promiscuity can be observed even between closely related pairs of alleles (e.g. at zero amino acid distance). As a result, there is no correlation between amino acid distance and promiscuity fold difference (Spearman’s rho = 0.02, P = 0.19). Amino acid distances were binned as shown on the figure (n = 308, 1168, 564, 654, 1492). Violin plots show the density function of promiscuity fold difference values for allele pairs in the given bin. White circles show median values, bold black lines show the interquartile range. (**B**) Sequence variability of an amino acid site in the epitope binding region of HLA-DRB1 (measured as Shannon entropy) correlates positively with the site’s promiscuity fragility, measured as the median predicted promiscuity fold difference caused by a random amino acid change at the given site (see inset, Spearman’s rho: 0.76, P = 0.0001). Sites that have a larger impact on promiscuity are more diverse in human populations. Line in inset represents linear regression between the two variables. The same result was obtained when promiscuity fragility was calculated based on nucleotide substitutions instead of amino acid substitutions (Spearman’s rho: 0.73, P = 0.0004, see Methods) or when sequence variability was measured as nonsynonymous nucleotide diversity (Π_*A*_), instead of sequence entropy (SI Appendix, Fig S7). Sites under positive selection as identified by Furlong et al.(30) show significantly higher promiscuity fragility (Wilcoxon rank sum test, P = 0.0012) and are marked with asterisks.

We next asked how selection on promiscuity has shaped the genetic diversity along the epitope binding region of HLA molecules. To quantify protein sequence variability at each amino acid position, we calculated the Shannon entropy index based on the alignment of the 96 selected HLA-DRB1 alleles from above. For each position, we also calculated promiscuity fragility, that is the median impact of single amino acid substitutions on promiscuity (See Methods). A strong positive correlation was found between Shannon entropy and promiscuity fragility (Fig. 4B, Spearman’s rho = 0.76, P = 0.0001). Importantly, this conclusion does not depend on how sequence polymorphism was estimated (SI Appendix, Fig. S7). Accordingly, amino acid positions with a large impact on epitope binding promiscuity are highly variable in human populations. Furthermore, those sites in the epitope binding region that are under positive selection (30) tend to have high promiscuity fragility values (Wilcoxon rank sum test, P = 0.0012, Fig. 4B). The above data suggests a link between allele promiscuity and HLA diversification, probably as an outcome of positive selection.

## Discussion

Central players of the adaptive immune system are the groups of proteins encoded in the major histocompatibility complex (MHC). By binding short peptide segments (epitopes), MHC molecules guide both immune response against pathogens and tolerance to self-peptides. The genomic region encoding these MHC molecules is of special interest, for two reasons. It harbors more disease associations than any other regions in the human genome, including associations to infectious diseases, autoimmune disorders, tumors and neuropsychiatric diseases (31). A growing body of literature is now revealing that certain MHC molecule variants can bind a wider range of epitopes than others, but the functional implications of this variation remain unknown (32). In this paper, we argue that by recognizing a larger variety of epitopes, such promiscuous MHC variants promote immune response against a broader range of pathogens at the individual level. Therefore, promiscuous epitope binding of MHC molecules should be favored by selection in geographic regions where extracellular pathogen diversity is high. Importantly, this mechanism is completely independent of the well-established concept of heterozygote advantage at the MHC, as it concerns individual alleles and not allele combinations or genotypes.

To test this hypothesis, we combined data on the geographic distribution of human MHC-II variants and prevalence of extracellular pathogens, empirical/computational estimates of epitope binding promiscuity and phylogenetic analyses. Our main findings, strongly supporting our hypothesis, are as follows.

First, in geographical regions of high extracellular pathogen diversity, human HLA DRB1 alleles have exceptionally high epitope binding repertoires. This suggests that the geographical distribution of promiscuous HLA-DRB1 alleles has been shaped by the diversity of extracellular pathogens. The HLA-DRB1*12:02 allele highlights this point. HLA-DRB1*12:02 is a promiscuous allele variant that has been associated with protection from certain infectious diseases. As expected, this allele is especially prevalent in regions of South East Asia with elevated parasite load (Fig. 3B). Notably, the relationship between pathogen diversity and epitope binding promiscuity may be more general as similar results hold for the HLA-A locus. HLA-A is one of the three major types of human MHC class I cell surface receptors, and is mainly involved in the presentation of epitopes from intracellular pathogens (33). In agreement with expectation, there is a strong positive correlation between local intracellular pathogen diversity and the HLA-A promiscuity level of the corresponding human populations (SI Appendix, Fig. S8, A and B, Dataset S2). No strong positive correlation was found for two other MHC class I genes (HLA-B and HLA-C, see SI Appendix, Fig. S8, C to F and Dataset S2). Therefore, other unrelated evolutionary forces may shape the geographical distribution of promiscuous HLA-B and HLA-C alleles (SI Appendix, Suppl. Text).

Second, a phylogenetic analysis revealed major differences in promiscuity levels of very closely related HLA-DRB1 alleles. This suggests that high promiscuity level in HLA-DRB1 has evolved rapidly and repeatedly during human evolution. Finally, amino acid positions with a prominent role in shaping HLA-DRB1 promiscuity level are especially variable in human populations and tend to be under positive selection. In sum, we conclude that HLA promiscuity level is a human trait with paramount importance during adaptation to local pathogens.

Our work has important ramifications for future studies. MHC is the most variable region of the human genome, and the variation is associated with numerous infectious and immune-mediated diseases (34–38). Pathogen-driven balancing selection is considered to generate the exceptionally high levels of genetic diversity, yet it cannot fully account for the observed geographic differences in human MHC genetic diversity (7, 25). Here we offer a complementary mechanism that provides protection against a broad range of pathogens at the individual level and shapes MHC genetic diversity at the same time. Therefore, promiscuity level may shape the global distribution pattern of human MHC II alleles.

We note that promiscuous MHC alleles are rare in certain human populations (SI Appendix, Table S1), suggesting that they are not always favoured by natural selection. Why should it be so? First, the advantage provided by individual promiscuous MHC alleles may be relatively small in regions with an exceptionally high MHC genetic diversity. Second, elevated promiscuity may not be able to cope with the rise of novel and highly virulent pathogens. In such cases, displaying a particular epitope might be the most efficient way to achieve resistance and high promiscuity might be suboptimal due to a reduced specificity (8). Third, high promiscuity level may elevate the risk of immune reactions against host tissues and non-harmful proteins (8, 39). Clearly, future work should elucidate the evolutionary trade-offs between protection from pathogens and genetic susceptibility to autoimmune diseases. This will require high-throughput experimental methods to determine epitope binding repertoire (40), and HLA transgenic mice studies on the role of promiscuity in immune response (41).

Finally, prior works indicate that genetic variations within particular MHC genes are known to influence vaccine efficacy (42), rejection rates of transplanted organs (43), susceptibility to autoimmune diseases (44) and antitumor immunity (45, 46). Our work raises the possibility that, by altering the maturation and functionality of the immune system, the size of the epitope binding repertoire of MHC variants itself could have an impact on these processes. The exact role of MHC promiscuity in these crucial public health issues is an exciting future research area.

## Methods

### Computational prediction of epitope-binding promiscuity

Epitopes of all available viral, bacterial and eukaryotic pathogens known to be bound by at least one HLA-I or HLA-II allele were collected from the Immune Epitope Database (IEDB) (21). Reference proteomes of pathogenic species that carry at least one of the collected epitope sequences were retrieved from the Uniprot database (102 for HLA-I and 71 for HLA-II epitopes) (47). Only epitopes of these species were analyzed further. All proteomes were scanned for each epitope sequence, and epitope sequences found in only one proteome (i.e. species-specific epitopes) were kept for further analysis. Highly similar epitope sequences were identified using Clustal Omega (48) and excluded as follows. A protein distance matrix was created and epitopes were discarded iteratively. In each iteration, the epitope pairs with the lowest k-tuple distance were identified. Then, the epitope with the highest average similarity to all other sequences was excluded. Iterations were repeated until distance values less than 0.5 (corresponding to greater than 50% sequence identity) were eliminated from the matrix (49). Note that this filtering procedure was carried out separately for epitope sequences bound by HLA-I and HLA-II.

Binding affinities of the remaining 3265 HLA-I epitope sequences to 346 HLA-A, 532 HLA-B and 225 HLA-C alleles were predicted with the NetMHCpan-4.0 algorithm. The binding of 2691 HLA-II epitope sequences to 606 HLA-DRB1 alleles was predicted with the NetMHCIIpan-3.1 algorithm (15). The “pep” sequence input format was used for both HLA-I and HLA-II epitope binding prediction. A binding affinity threshold of 50 nM was applied, yielding peptides that are likely to be immunodominant (22). For alternative binding threshold definitions, see SI Appendix, Fig. S4. For each binding threshold, epitope-binding promiscuity was defined as the fraction of the epitope set bound by a given allele.

### Calculating epitope-binding promiscuity using in vitro data

To determine the epitope binding promiscuity of HLA-DRB1 alleles based on previously published experimental data, we used the IEDB database (21). Specifically, we downloaded all MHC ligand and T cell assay data available for 48 HLA-DRB1 alleles. Binding data of 20 alleles screened for at least 100 ligands were further analyzed. The epitope set of each allele was filtered for highly similar sequences as described above. As the majority of in vitro assay data were available in a binary format (i.e. presence or absence of binding), promiscuity was calculated as the fraction of positive binding assays for a given allele.

### Calculating promiscuity levels of human populations

To calculate population-level promiscuity values, we obtained HLA allele frequency data from the Allele Frequency Net Database (AFND) and the International Histocompatibility Working Group (IHWG) populations (18, 19). Haplotype-level data of the 13^*th*^ International HLA and Immunogenetics Workshop (IHIW) populations were downloaded from dbMHC (National Center for Biotechnology Information [NCBI]; http://www.ncbi.nlm.nih.gov/mhc/). Additionally, allele frequency data of the 14th and 16th IHIW populations as published by Riccio et al. (20) and populations in the AFND were used in the analyses. To avoid potential confounding effects of recent genetic admixture and migration, we focused on native populations, similarly to previous studies (SI Appendix, Table S1) (7, 25). We excluded IHWG populations reported to deviate from Hardy-Weinberg equilibrium (20). Among the AFND populations and IHWG populations without haplotype-resolution data (14th and 16th IHIW), those comprising less than 100 genotyped individuals or those in which the sum of allele frequencies deviated from 1 by more than 1% were excluded. Populations reported in multiple databases were included only once in the analysis.

For each HLA loci, we calculated mean population promiscuity by averaging promiscuity values of alleles weighted by their relative frequencies in the populations. In all of these calculations, we used standardized (i.e. z-score) promiscuity values to make the in silico and in vitro values more easily comparable. Finally, when calculating population-level promiscuity based on in vitro promiscuity data, we excluded populations for which in vitro promiscuity values could be assigned to less than 50% cumulative allele frequency.

To tackle the issue of non-independence of data points, we focused on populations instead of countries and grouped those populations that have highly similar HLA allele compositions, based on standard measures of genetic distance (see below). We merged populations with highly similar HLA allele compositions, allowing us to avoid pseudoreplication of data points while retaining informative allele frequency differences between populations residing in the same broad geographical areas.

To this end, we first generated a genetic distance matrix between populations with the adegenet R library using allele frequency data of the examined locus. We used the Rogers’ genetic distance measure (50) because it doesn’t assume that allele frequency changes are driven by genetic drift only, an unlikely scenario for HLA genes. Next, populations were merged using a network-based approach. Populations were treated as nodes and two nodes were connected if their genetic distance was under a cutoff value. Populations were grouped in an iterative manner. In each iteration, all maximal cliques (i.e. subsets of nodes that are fully connected to each other) in the network were identified. Maximal cliques represent groups of populations where all populations have similar allele compositions to each other. Then, mean genetic distance within each clique was calculated. The clique with the lowest average distance was selected and its populations were grouped together. Then, this clique was deleted from the network. Iterations were repeated until no maximal cliques remained in the network. Grouping of populations was carried out using different distance value cutoffs (1^*st*^, 5^*th*^, 10^*th*^ and 15^*th*^ rank percentile of all distance values). The resulting population groups and the individual populations remained in the network were treated as independent data points in subsequent statistical analyses. Mean promiscuity level in population groups was calculated by averaging population promiscuity values.

Unless otherwise indicated, all figures are based on population groups using the 15^*th*^ percentile genetic distance cutoff value. Importantly, using different cut-offs has no impact on our results (Dataset S2). Finally, we note that genetic differences among human populations mostly come from gradations in allele frequencies rather than from the presence of distinctive alleles (51). Therefore, traditional clustering of populations based on HLA composition would have been ill-suited for our purposes, yielding only a small number (3-4) of clearly distinct clusters (data not shown).

### Pathogen diversity

Data on 309 infectious diseases were collected from the Global Infectious Diseases and Epidemiology Network (GIDEON) (24). For each disease, the number of causative species or genera (when species were not listed for the genus) was determined using disease information in the GIDEON database as described previously (52). Causative agents were classified into obligate intracellular and extracellular pathogen groups based on a previous study (7) and literature information. Putative facultative intracellular pathogens were excluded from the analysis. Diseases caused by agents that could not be clearly classified were also excluded from the analysis. Extracellular and intracellular pathogen diversity (richness) of each country was approximated by the number of identified endemic extracellular and intracellular species, respectively.

Finally, we assigned country-level measures of pathogen and HLA diversity to population groups as follows. For each population group, extracellular and intracellular pathogen counts were calculated by averaging the corresponding country-level values across the populations within the group. For example, if a population group contained two populations residing in neighboring countries then we assigned the average pathogen diversity of the two countries to it.

### Amino acid distance between DRB1 alleles

We used amino acid distance as a proxy for phylogenetic distance between pairs of DRB1 alleles. To this end, nucleotide sequences of DRB1 alleles that contained full exon 2 and 3 regions were downloaded from the IPD-IMGT/HLA database (10). To limit our analyses to alleles that have an impact on the inferred promiscuity level of a population, we considered only those sequences that had a non-zero frequency in at least one human population (see above). From allele groups that code for the same protein sequence (synonymous differences, differentiated by the third set of digits in the HLA nomenclature), one of the alleles was randomly chosen. This selection procedure resulted in 96 alleles. Multiple alignment of nucleotide sequences was performed using the MUSCLE algorithm as implemented in the MEGA software (53) and converted to protein sequence alignments. Amino acid distance was calculated using the Jones-Taylor-Thornton substitution model in MEGA (53) (Fig. 4A). Epitope binding region sites - as defined previously (15) - were excluded when calculating amino acid distance. The rationale behind this exclusion is that these sites are known to be under positive selection (54, 55) and are therefore less informative on evolutionary distance. Additionally, by removing these sites, the amino acid distance remains independent of promiscuity predictions. Finally, as intragenic recombination may distort the inference of evolutionary distance, we identified such events across all alleles following the protocol of Satta et al. (56) using GENECONV (57) and RDP algorithms (58) as implemented in the RDP software (59). Recombinant alleles were removed when calculating amino acid distance.

### Sequence diversity and promiscuity fragility

We first defined the epitope binding region of HLA DRB1 alleles as previously (15). To estimate sequence diversity along the epitope binding region, we employed two measures: standard Shannon entropy (60) and nucleotide diversity (Π), a widely employed measure of genetic variation (61).

Using the protein sequence alignment of the 96 alleles defined above, we calculated amino acid sequence variability as the Shannon entropy of the given amino acid site as follows:

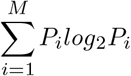

where *P_i_* is the fraction of residues of amino acid type i at a given site and M is the number of amino acid types observed at that site.

Nonsynonymous nucleotide diversity (Π_*A*_) measures the average number of nonsynonymous nucleotide differences per nonsynonymous site between two randomly chosen protein coding DNA sequences from the same population (61, 62). Π_*A*_ was calculated for each amino acid site in the epitope binding region for each population using DnaSP software (63) and custom-written R scripts. Nucleotide sequences of DRB1 alleles were downloaded from the IPD-IMGT/HLA database (10).

The calculation is based on the work of Nei et al. (61) using the equation:

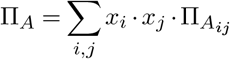

where *x_i_* and *x_j_* are the frequencies of the ith and jth alleles in the population, respectively and 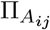 is the number of nonsynonymous nucleotide differences per nonsynonymous nucleotide site between the two codon sequences of the given amino acid site in the *i^th^* and *j^th^* alleles. To calculate Π_*A*_ for each population, allele frequency data of human populations was obtained, as described earlier (see above). An overall nucleotide diversity index was calculated by averaging Π_*A*_ across populations.

To calculate each amino acid site’s impact on epitope binding promiscuity (promiscuity fragility), promiscuity was predicted for each 19 possible amino acid change along the epitope binding region of each 96 alleles. The fold difference in promiscuity resulting from each amino acid substitution was calculated. The median promiscuity fold difference of each possible allele and amino acid change combination (96 × 19) was used to estimate promiscuity fragility at each amino acid position. As some of the 19 possible amino acid changes are not accessible via a single nucleotide mutation and the accessible amino acid changes can have different likelihoods based on the codon sequence of the site and the genetic code, we also calculated promiscuity fragility based on each non-synonymous nucleotide substitution of the codon instead of each amino acid substitution of the site.

### Statistical analysis and graphical representation

All statistical analyses were carried out in R version 3.2.0 (64). Smooth curves were fitted using the cubic smoothing spline method (65).

## ACKNOWLEDGEMENTS

We wish to thank Jim Kaufman for the insightful comments we have received on an earlier version of the manuscript. We are also grateful to Károly Kovács for his suggestions on data analysis. The authors acknowledge the following financial support: “Frontline” Research Excellence Programme of the National Research, Development and Innovation Office, Hungary (KKP_17), “Lendület” Programme of the Hungarian Academy of Sciences and The Wellcome Trust (BP and CP), H2020 ERC-2014-CoG (CP), GINOP-2.3.2-15-2016-00014 (EVOMER, BP and CP), GINOP-2.3.2-15-2016-00020 (MolMedEx TUMORDNS, CP) and GINOP-2.3.3-15-2016-00001 (CP), GINOP-2.3.2-15-2016-00057 (“Az evolúció fényében: elvek és megoldások”, VM) German Research Foundation grant LE 2593/3-1 (TLL). Máté Manczinger was supported by the UNKP-17-4 New National Excellence Program of the Ministry of Human Capacities; Viktor Müller was supported by a Bolyai János Research Fellowship of the Hungarian Academy of Sciences.

